# ATF4 and EcR interact to mediate both transcriptional activation and repression in the *Drosophila* fat body

**DOI:** 10.64898/2026.03.10.710866

**Authors:** Manuel H. Michaca, Lydia Grmai, Deepika Vasudevan

**Author notes:** To whom correspondence must be addressed.

## Abstract

Cells with high secretory and metabolic loads such as adipocytes and hepatocytes rely on constitutive activation of stress response pathways for their homeostatic function and to cope with exogenous stressors such as nutrient deprivation or excess lipids. The evolutionarily conserved stress response factor, Activating Transcription Factor 4 (ATF4), is known to be required for both homeostatic function and exogenous burden in these tissues. However, the molecular mechanism by which ATF4 specifies distinct transcriptional targets under homeostasis versus stress conditions remains an open question. Here, we use the *Drosophila* larval fat tissue as a model to establish that ATF4 interacts with the steroid hormone receptor Ecdysone Receptor (EcR) to transcriptionally activate and repress genes involved in lipid metabolism. Our data show that EcR and its ligand, 20-hydroxyecdysone, are required for transcription activation of the *bona fide* ATF4 target *4E-BP* in the fat body. We also find that ATF4 and EcR co-repress transcription of the triglyceride lipase *brummer* (*bmm*). In Förster resonance energy transfer (FRET) experiments, we find that ATF4 interacts with EcR and does so by competing with the canonical EcR-binding partner, Ultraspiracle (Usp). Using a genetic model of nutrient deprivation, we find that while EcR is required for homeostatic signaling, it is dispensable for the elevated ATF4 signaling associated with nutrient deprivation as measured by induction of *4E-BP*. Together, these data provide a mechanistic starting point for understanding how changes in interaction partner allows ATF4 to engage in context-specific transcriptional programs in metabolic tissues.

## Introduction

Cells have evolved stress response mechanisms to cope with a variety of insults, such as nutrient deprivation, protein misfolding, chemical insults, and others. Certain tissue types in multi-cellular organisms, due to the nature of their function, have co-opted these mechanisms for their homeostatic function. Classic examples of such function include high secretory demand, constant metabolic turnover, and toxin exposure^1–5^. Notably, metabolically active tissues, such as adipose and liver, rely on constitutive activation of the Integrated Stress Response (ISR) pathway for their homeostatic function^1,6–9^. In addition to this homeostatic activity in metabolic tissues, aberrantly high ISR activation has been associated with a variety of disorders including hyperlipidemia and steatohepatitis^10–12^. While there are many stress response signaling pathways that are activated by these disorders, the defining feature of ISR signaling is its duality: it supports homeostatic function of metabolic cell types, but its hyperactivation due to further exogenous stress can have deleterious consequences^6,7,11,13–17^. However, the molecular distinctions between homeostatic and stress functions of ISR signaling, particularly in adipocytes, are not well understood.

The ISR is a highly conserved signaling pathway that detects stress via kinases that enable selective translation of the master transcription factor Activating Transcription Factor 4 (ATF4)^18–20^. ATF4 is a member of the basic leucine zipper (bZIP) transcription factor family. Leucine zipper domains typically allow for stable DNA binding through homo- or heterodimerization. The leucine zipper is also thought to define binding partner specificity^21,22^. Notably, the ATF4 leucine zipper domain has been structurally demonstrated to be unsuited for homodimerization, and this is also supported by *in vitro* binding studies^23–25^. Previous bZIP network analyses have predicted several heterodimeric interaction partners for ATF4, including CCAAT/enhancer-binding proteins α-ε (C/EBPα-ε), C/EBP homologous protein (CHOP), cAMP-responsive element binding protein 3 like 2 (CREB3L2), X-box binding protein 1 (XBP1), ATF2, ATF3, Musculoaponeurotic Fibrosarcoma factor (MAF), JUN, and FOS^23,24,26^. Many of these interactions have been functionally validated *in vivo* and *in vitro*^27^. Of the known interaction partners of ATF4, C/EBPβ is the best studied in adipocytes. The ATF4-C/EBPβ complex has been demonstrated to be required for adipocyte differentiation^28,29^, though its role in homeostatic adipocyte function or in specific disease states is not known.

ATF4 is an important regulator of lipid metabolism in adipose tissues^1,6–9^. *ATF4* null mutant mice have decreased overall triglycerides and changes in the expression of several lipid metabolism genes including adipocyte triglyceride lipase (ATGL) and several perilipins^6,7^. Similar lean phenotypes have been observed in *Drosophila*, where loss-of-function mutants of ATF4 (encoded by *cryptocephal, crc*) show reduced overall triglyceride levels and increased susceptibility to nutrient deprivation^6,30^. In *Drosophila*, ATF4 is basally active in adipose tissue (called the fat body) during both larval and adult stages and is required for its homeostatic function during both stages^1,8,31,32^. Interestingly, ATF4 has also been implicated in regulating molting behavior during larval development, but the exact tissue type where ATF4 is required for such regulation is unknown^33,34^. As with vertebrate ATF4, several bZIP interaction partners have been put forth for *Drosophila* ATF4 based on *in vitro* experiments^35^. *Drosophila* ATF4 has also been shown to directly interact *in vitro* with a non-bZIP transcription factor, Ecdysone receptor (EcR)^34^. While non-bZIP interaction partners have also been proposed for vertebrate ATF4, such as Runx2^36,37^, there are no reports on physiological roles for ATF4-EcR interaction to the best of our knowledge.

EcR is a nuclear hormone receptor that is best known for its role in developmental progression in insects^38,39^. EcR activates transcriptional targets when bound to the steroid 20-hydroxyecdysone (20E), the levels of which are tightly regulated across development^40–42^. *Drosophila* EcR has three splice isoforms that result in protein products with distinct N-terminal regions but identical ligand binding and DNA-binding regions: EcR-A, EcR-B1, and EcR-B2^43^. EcR is known to canonically interact with Ultraspiracle (Usp) to mediate EcR transcriptional activity^44^. While EcR is best studied in the context of developmental timing, it is also known to regulate lipid metabolism and respond to stress^10–12,45,46^. Interestingly, ATF4 and EcR-B2 were previously identified as interaction partners in a yeast two-hybrid screen and the interaction was confirmed using *in vitro* binding assays^34^. However, whether this putative interaction between ATF4 and EcR occurs *in vivo*, which cell types utilize such interaction, and the biological role of cooperative ATF4-EcR function all remain unknown. In this study, we use the *Drosophila* larval adipose tissue to test whether ATF4 interacts with EcR in developing adipocytes and whether this changes with exogenous stress. Our work presents a mechanistic understanding of how metabolically active tissues with constitutive ATF4 signaling engage different transcriptional outputs under homeostasis and stress conditions.

## Results

### EcR interacts with ATF4 to regulate its transcriptional activity in the fat body

We and others have shown that the developing third instar larval fat body shows high levels of ATF4 activity^47,48^. Since ATF4 has previously been shown to interact with EcR *in vitro*, we asked whether EcR is required for ATF4 activity in the fat body^34^. We first tested whether loss of EcR resulted in changes in *4E-BP* expression (also known as *Drosophila Thor*)^49^, a known transcriptional target of ATF4 in the larval fat body^48,50^. We knocked down *EcR* using a fat body driver, *Dcg-GAL4* ^51^, and performed RT-qPCR on isolated fat bodies. We found that *Dcg>EcR^RNAi^* fat bodies showed a robust decrease in *4E-BP* transcript levels, to an even greater extent than *Dcg>ATF4^RNAi^*fat bodies (**Fig. 1A**). Notably, our analysis showed that RNAi-mediated knockdowns were effective, as seen by a reduction in *ATF4* and *EcR* transcript levels in *Dcg>ATF4^RNAi^* and *Dcg>EcR^RNAi^*animals, respectively. Further, our RT-qPCR analyses showed that depleting *EcR* does not affect *ATF4* transcript levels (**Fig. 1A**), supporting our premise that EcR regulates ATF4 transcriptional activity and not through depletion of ATF4.

**Fig 1.**
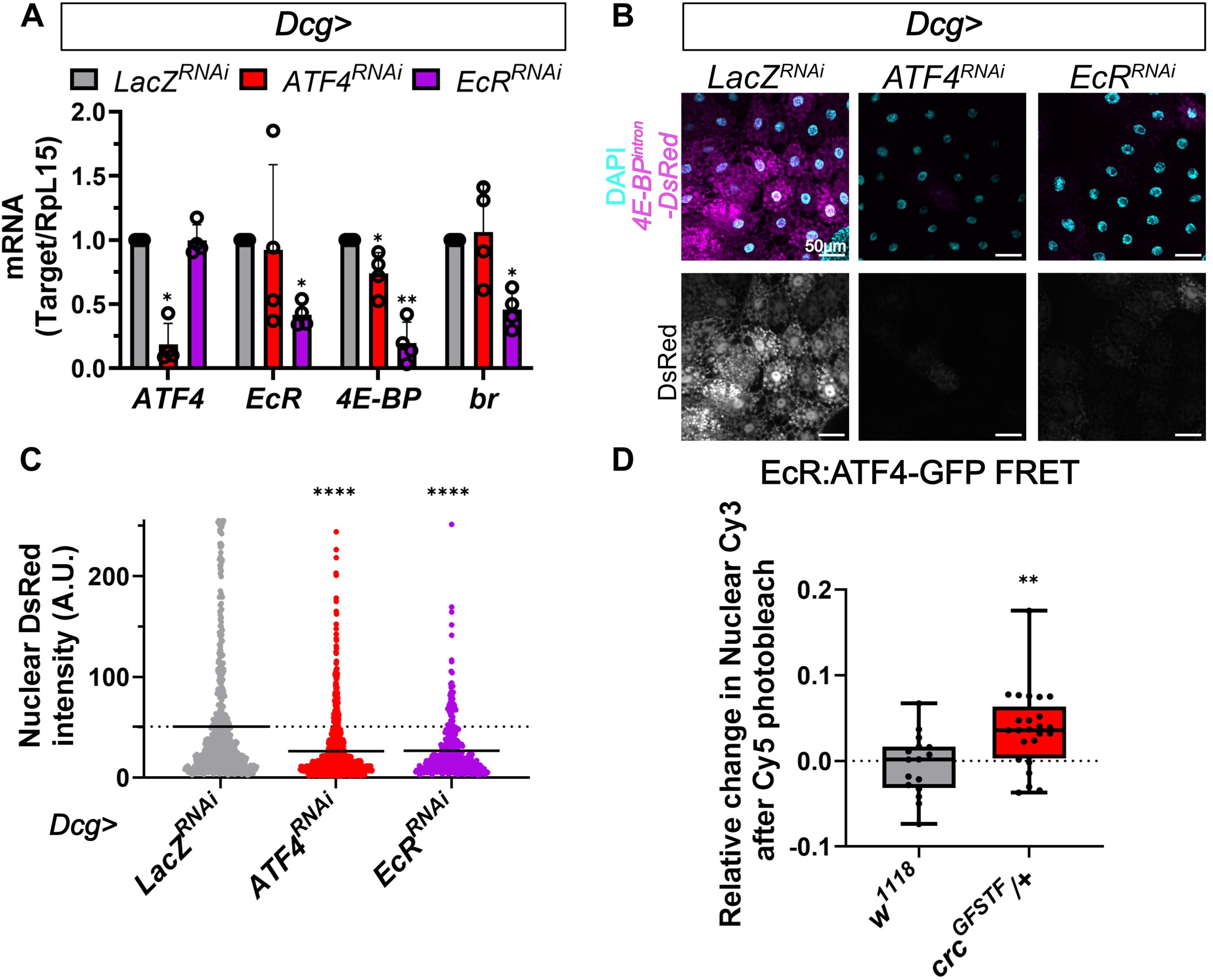
EcR regulates ATF4 transcriptional activity in the wandering third instar larval fat body. **A)** RT-qPCR analysis of indicated transcripts in isolated fat bodies from female animals expressing *UAS* transgenes encoding control (*LacZ^RNAi^*), *ATF4^RNAi^*or *EcR^RNAi^*. *UAS*-transgene expression is driven by *Dcg-GAL4*. Data represent average of four biological replicates and error bars represent standard error of mean. **B)** Representative confocal images showing 4E-BP^intron^-DsRed reporter expression (magenta) in female fat bodies also expressing indicated RNAi lines driven by *Dcg-GAL4*. Nuclei are counterstained with DAPI (cyan). **C)** Quantification of nuclear DsRed in individual adipocytes from **B**. Data represent mean from at least 18 animals collected from three independent crosses. The dotted line indicates the average of the control (*LacZ^RNAi^*) sample. **D)** Relative Change in FRET after Acceptor (ATF4-GFP labeled with Cy5) photobleach upon excitation of the donor (EcR labeled with Cy3) in the nuclei of adipocytes from control (*w^1118^*) or ATF4-GFP expressing (*crc^GFSTF^/+*) female animals. Data represent mean from at least 18 animals collected from three independent crosses. The dotted line indicates the average of the control (*w^1118^*) sample. Scale bar=50 μm. ****=p<0.00001; ***=p<0.0001; **=p<0.001; *=p<0.05. Only significant comparisons are shown. Statistical tests are described in the methods.

Next, we sought to test whether EcR specifically regulated *4E-BP* via the ATF4-binding sites in the *4E-BP* locus. To do so, we utilized the *4E-BP^intron^-DsRed* transgenic reporter^48^, where the intronic region of *4E-BP* containing ATF4 binding sites is placed upstream of *DsRed* and serves as a highly specific readout of ATF4 activity. As expected, *Dcg>ATF4^RNAi^* animals showed a marked decrease in nuclear DsRed intensity in comparison to control fat bodies from *Dcg>LacZ^RNAi^* animals (**Fig. 1B, C**). We observed a similar decrease in DsRed intensity in *Dcg>EcR^RNAi^* fat bodies (**Fig. 1B, C**), which we also corroborated using another independent *EcR^RNAi^* line (**Fig. S1A-B**). Since we have previously reported that 4E-BP^intron^-DsRed levels increase in the fat body between second and third instar larval stages^48^, we sought to exclude the possibility that the reduced DsRed intensity seen in *Dcg>EcR^RNAi^*fat bodies could be due a developmental delay in these animals. To test this, we measured nuclear size across all genotypes in (**Fig. 1B**) since adipocyte cell size and volume has been demonstrated to increase between second and third instar larval stages^52^. We found that nuclear size from *Dcg>ATF4^RNAi^* and *Dcg>EcR^RNAi^*fat bodies were not decreased in comparison to control nuclei from *Dcg>LacZ^RNAi^*(**Fig. S1C**). These data assured us that the differences observed from 4E-BP^intron^-DsRed in (**Fig. 1B, C**) cannot be ascribed to a developmental delay but are due to EcR-mediated regulation of ATF4 transcriptional activity. Further, though we observed such differences in both female (**Fig. 1B-C**) and male animals (**Fig. S1A-B**), we chose to focus on females for the rest of this study for ease of comparison.

Given that EcR has been previously shown to bind to ATF4 *in vitro*^34^, we considered whether EcR regulation of ATF4 transcriptional activity in the fat body relies on interaction between these two transcription factors. To detect *in vivo* interaction between ATF4 and EcR, we performed a fluorescence resonance energy transfer (FRET) assay on immunolabeled ATF4 and EcR, using selective photobleaching to examine FRET changes. We immunolabeled endogenous EcR with a donor fluorophore (Cy3). Additionally, we utilized MiMIC^53^-replacement transgenic animals wherein endogenous ATF4 is tagged with GFP (*crc^GFSTF^*)^32,53,54^ and labeled GFP with an acceptor fluor (Cy5). Since Cy5 is excited in the Cy3 emission range, the quantifiable Cy3 emission is lower when the two fluors are proximal to each other. In comparison to *w^1118^* controls lacking ATF4-GFP, fat bodies from *crc^GFSTF^/+* expressing one copy of ATF4-GFP-Cy5 showed significant relative change in FRET after photobleaching of EcR-Cy3 (**Fig. 1D**). Since energy transfer between occurs only if the compatible fluorophores are within 10nm of each other^55^, these data demonstrate that indeed ATF4 and EcR directly interact with each other *in vivo*.

### ATF4 largely does not affect EcR transcriptional activity in the fat body

Since our data find that EcR regulates ATF4 transcriptional activity, we asked whether ATF4 likewise regulates EcR transcriptional activity. We first tested this by measuring transcript levels of *broad* (*br*), an EcR target^56^. As expected, we observed a decrease in *br* in *Dcg>EcR^RNAi^*animals using RT-qPCR (**Fig. 1A**). However, *Dcg>ATF4^RNAi^* fat bodies showed no change in *br* levels (**Fig. 1A**), suggesting that ATF4 may not be required for transcription of EcR target genes in the fat body. To more comprehensively test the ATF4-EcR regulatory paradigm, we performed transcriptomic analysis (RNAseq) on fat bodies isolated from control (*Dcg>LacZ^RNAi^*), *Dcg>ATF4^RNAi^*, and *Dcg>EcR^RNAi^* wandering third instar larvae. Consistent with our qPCR results in **Fig. 1A**, RNAseq data revealed that depleting ATF4 did not impact *EcR* transcript levels and vice versa (**Fig. S2**). Loss of *ATF4* and *EcR* individually resulted in differential expression of 1378 and 1841 genes, respectively (**Fig. 2A**). Loss of *ATF4* resulted in downregulation of 934 transcripts, 308 of which were downregulated upon loss of EcR, and in the upregulation of 444 transcripts, 216 of which were upregulated upon loss of EcR (**Fig. 2A-B**).

**Fig. 2.**
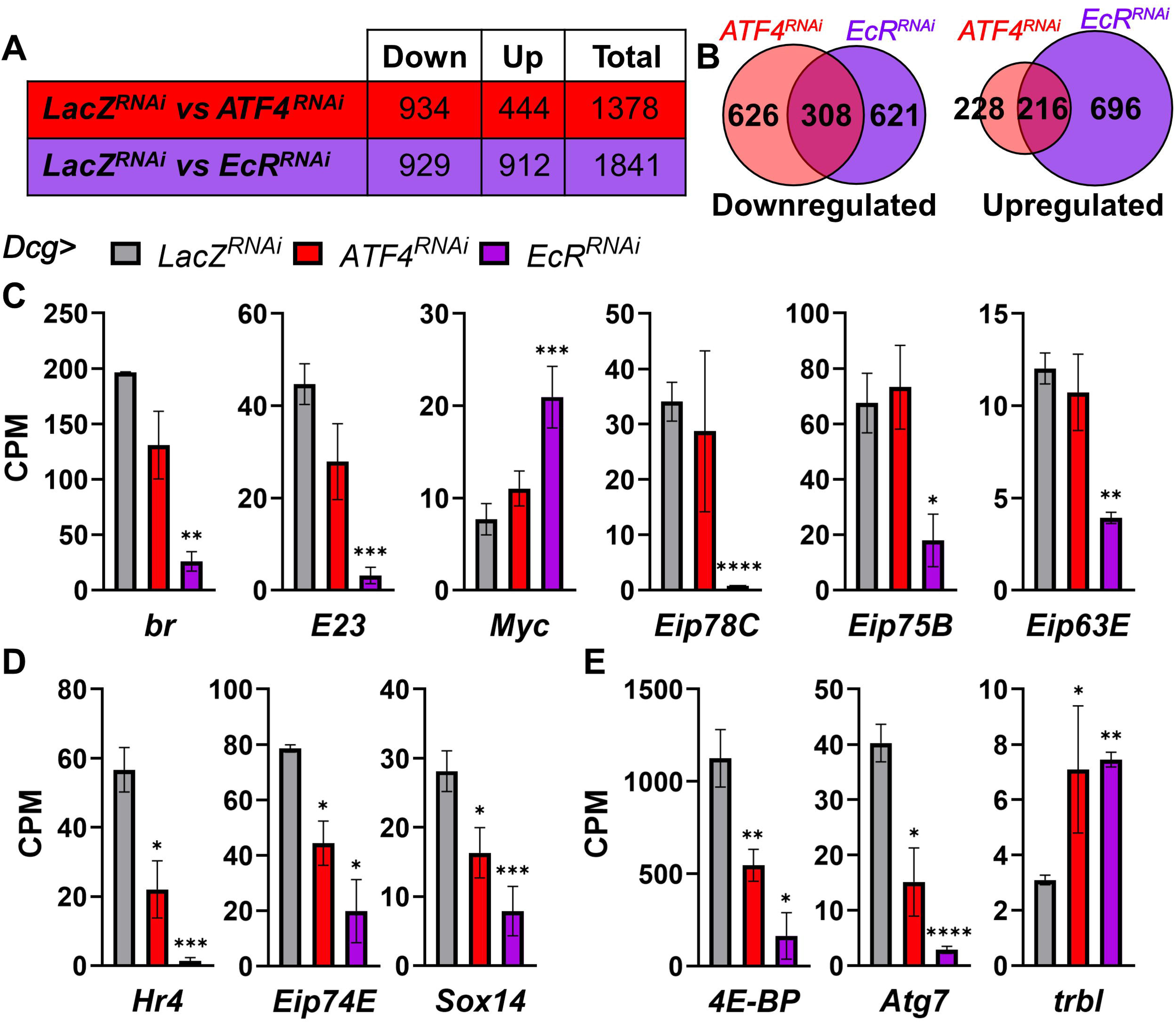
ATF4 depletion largely does not impact EcR transcriptional activity. **A)** Table summarizing the overall count of differentially regulated genes in RNAseq analyses performed in isolated fat bodies where the indicated *RNAi* lines are driven by *Dcg-GAL4*. **B)** Venn-diagram showing the commonalities in the differentially downregulated (left) and upregulated genes (right) in **A.** **C-E)** Counts per million (CPM) for the indicated transcripts from the RNAseq analysis in **A**. **C** and **D** show data for known EcR targets, **E** shows data for known ATF4 targets. Also see **Table S1** and **S2**. ****=p<0.00001; ***=p<0.0001; **=p<0.001; *=p<0.05. Only significant comparisons are shown. Statistical tests are described in the methods.

We examined changes in known EcR targets; *br*, *E23*, *Myc*, *Sox14*, *Hr4*, and the ecdysone induced proteins (Eips) *Eip78C*, *Eip75B*, *Eip63E*, and *Eip74EF*^57,58^ in our RNAseq data sets (**Tables S1**, **S2**). While all these targets were differentially regulated in *Dcg>EcR^RNAi^* (**Fig. 2C-D**), many of these targets were unchanged in *Dcg>ATF4^RNAi^*(**Fig. 2C**). There were a few exceptions, namely *Hr4*, *Eip74EF*, and *Sox14*, which showed a modest change with the loss of ATF4 (**Fig. 2D**). We also examined changes in the transcription of known ATF4 targets found to be differentially expressed by RNAseq analysis (see **Tables S1** and **S2)**, including *Thor*, *Atg7*, and *trbl*^27,48^. Per our hypothesis that EcR is required for ATF4 transcriptional activity, all these targets were differentially regulated in *Dcg>EcR^RNAi^* (**Fig. 2E**). Given that the majority of EcR targets were unchanged with ATF4 depletion, we conclude based on these analyses that while EcR is required for ATF4 transcriptional activity, ATF4 largely does not impact EcR transcriptional activity in the third instar larval fat body.

### ATF4 activity in larval adipocytes relies on 20-hydroxyecdysone

We next sought to test whether known components of EcR signaling also regulated ATF4 activity in the fat body. Canonically, EcR activates target gene transcription upon binding to 20-hydroxyecdysone (20E) and a host of interaction partners, the best studied of which is the transcription factor Usp^44,59^. In larvae, the steroid hormone ecdysone is produced in the ring gland and released at specific developmental time points which correspond to initiation of the various molts and metamorphosis^60^. Circulating ecdysone has bene demonstrated to be actively taken up by Ecdysone importer (EcI) in most cell types, including adipocytes^61^. Imported ecdysone is converted to the active form 20E by the cytochrome P450 monooxygenase, Shade (Shd)^62^. We began by testing whether depletion of *EcI or shd*, either of which would result in decreased 20E levels, impacts ATF4 activity in the fat body as measured by *4E-BP^intron^-DsRed*. We found that both *Dcg>EcI^RNAi^* and *Dcg>shd^RNAi^* resulted in a statistically significant decrease in DsRed levels (**Fig. 3A, B**). As previously, we confirmed that our experiments compared animals at similar developmental stages by measuring nuclear size (**Fig. S1C**). These data together indicate that 20E is required for ATF4 activity in the fat body.

**Fig 3.**
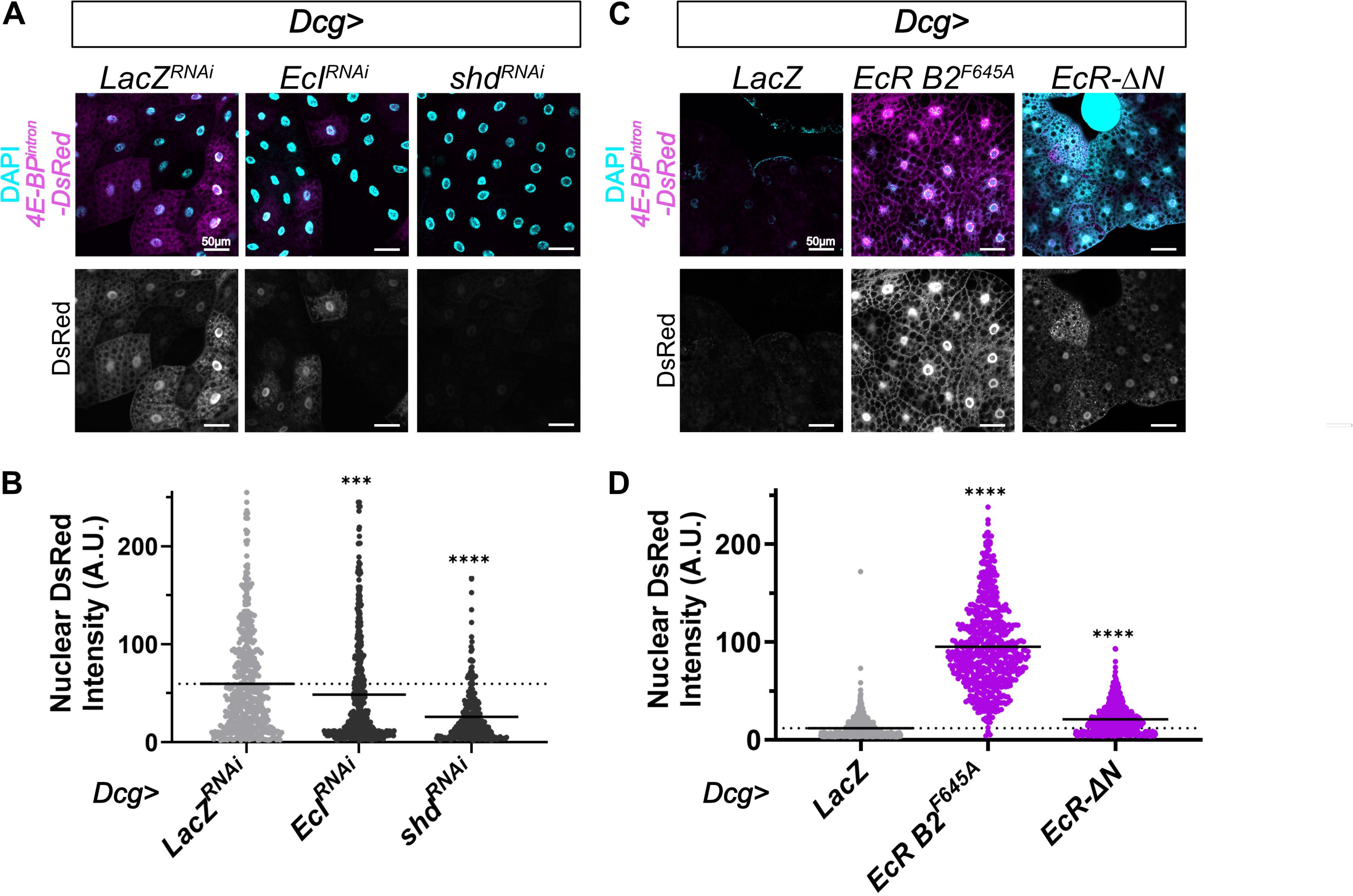
20E is required for ATF4 Transcriptional activity. **A)** Representative confocal images showing 4E-BP^intron^-DsRed reporter expression (magenta) in fat bodies also expressing *LacZ^RNAi^*(control), *EcI^RNAi^* or *shd^RNAi^* driven by *Dcg-GAL4*. **B)** Quantification of nuclear DsRed in individual adipocytes from **A**. Data represent mean from at least 18 animals collected from three independent crosses. The dotted line indicates the average of the control (*LacZ^RNAi^*) sample. **C)** Representative confocal images showing 4E-BP^intron^-DsRed reporter expression (magenta) in fat bodies also expressing control (*LacZ*), *EcR^F654A^,* or *EcR-ΔN* driven by *Dcg-GAL4*. **D)** Quantification of nuclear DsRed in individual adipocytes from **C**. Data represent mean from at least 18 animals collected from three independent crosses. The dotted line indicates the average of the control (*LacZ*) sample. Scale bar=50 μm. ****=p<0.00001; ***=p<0.0001; **=p<0.001; *=p<0.05. Only significant comparisons are shown. Statistical tests are described in the methods.

While all three EcR isoforms (-A, -B1, and -B2) are known to bind 20E and Usp through the ligand-binding domain, *in vitro* experiments show that only EcR-B2 can interact with ATF4 via its N-terminal region^34,43^. Based on this, we posited that expressing a transgenic EcR lacking the N-terminal domain (*EcR-*Δ*N*) would not impact ATF4 transcriptional activity in the fat body. Indeed *Dcg>EcR-*Δ*N* showed only a small, albeit statistically significant, effect on *4E-BP^Intron^-DsRed* (**Fig. 3C-D**). In contrast, expression of a full length, dominant-negative EcR-B2 (*Dcg>EcR-B2^F645A^*) resulted in a striking increase in *4E-BP^Intron^-DsRed* (**Fig. 3C-D**). Notably, the point mutation in this EcR variant (phenylalanine-to-alanine replacement) does not disrupt interaction with 20E or Usp. However, the mutation does render the variant inactive as determined using EcR reporters^63–65^. Though EcR-B2^F645A^ has a dominant negative effect on EcR signaling^63–65^, our data show that expressing EcR-B2^F645A^ increases ATF4 activity (**Fig. 3C-D**). Thus, we propose that the transcriptional activity of the putative ATF4-EcR complex is mechanistically distinct from that of the EcR-Usp complex. Further, since we observed that increasing EcR availability by overexpressing EcR-B2^F645A^ increases ATF4 activity, we predict that EcR levels in adipocytes are limited and this conjecture is experimentally tested below.

### ATF4 and Usp compete for EcR in adipocytes

Since our analyses indicated that 20E is required for EcR activity, we next asked whether the canonical EcR-binding partner, Usp, is required for ATF4 transcriptional activity^65,66^. In contrast to our results with *shd^RNAi^* and *EcI^RNAi^*(**Fig. 3A-B**), we found that *Dcg>usp^RNAi^*fat bodies showed an increase in *4E-BP^intron^-GFP* reporter expression (**Fig. 4A-B**), which is the GFP variant of the *4E-BP^intron^-DsRed* reporter^30^. This led us to consider that depletion of *usp* resulted in increased ATF4 activity due to increased EcR availability, particularly since we previously surmised that EcR levels in adipocytes may be limiting. To ensure this increase was indeed due to increased ATF4 activity, we simultaneously knocked down both *ATF4* and *usp*. This resulted in diminished *4E-BP^intron^-GFP* expression in comparison to controls, abolishing the effect of *usp* knockdown on reporter activity (**Fig. 4A-B**). We note that the primary reason we utilized the GFP variant of the *4E-BP^intron^*reporter in these experiments is to facilitate ease of semi-automated quantification of nuclear GFP as reported previously^67^. These data led us to posit that Usp likely competes with ATF4 for binding with EcR, such that depletion of *usp* results in increased EcR availability for ATF4-EcR complex formation and thus increased ATF4 activity. We tested this hypothesis in FRET experiments as described below.

**Fig. 4.**
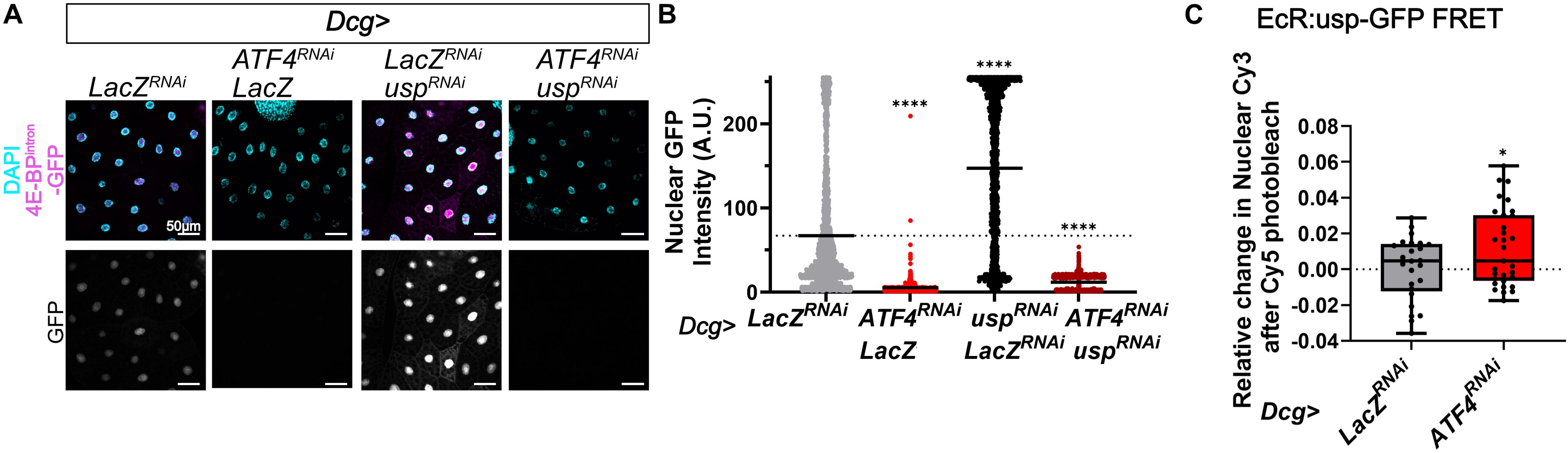
ATF4 competes with Usp to interact with EcR. **A)** Representative confocal images showing 4E-BP^intron^-GFP reporter expression (magenta) in fat bodies also expressing *LacZ^RNAi^*, either *ATF4^RNAi^*or *usp^RNAi^,* and both *usp^RNAi^* and *ATF4^RNAi^*driven by *Dcg-GAL4.* Equal number of UAS-transgene cassettes were used by combining RNAi lines with either *LacZ* or *LacZ^RNAi^*. **B)** Quantification of nuclear GFP in individual adipocytes from **A**. Data represent mean from at least 18 animals collected from three independent crosses. The dotted line indicates the average of the control (*LacZ^RNAi^*) sample. **C)** Relative Change in FRET after Acceptor (Usp-GFP labeled with Cy5) photobleach upon excitation of the donor (EcR labeled with Cy3) in the nuclei of wandering third instar larval fat bodies where *Dcg-GAL4* drives expression of control (*LacZ^RNAi^*) or *ATF4^RNAi^*. Data represent mean from at least 18 animals collected from three independent crosses. The dotted line indicates the average of the control (*LacZ^RNAi^*) sample. Scale bar=50 μm. ****=p<0.00001; ***=p<0.0001; **=p<0.001; *=p<0.05. Only significant comparisons are shown. Statistical tests are described in the methods.

Based on our hypothesis that ATF4 and Usp compete for binding to EcR in adipocytes, we predicted that depleting ATF4 should result in increased EcR-Usp interaction. To test this, we designed a FRET experiment similar to **Fig. 1D** wherein we immunolabeled endogenous EcR with a donor fluor (Cy3) and utilized a transgenic Usp-GFP construct (*PBac{usp-GFP}*) immunolabeled with an acceptor fluor (Cy5). We measured the relative change in FRET after acceptor photobleaching in control (*Dcg>LacZ^RNAi^*) fat bodies and *Dcg>ATF4^RNAi^*. Consistent with our hypothesis, we found that depleting ATF4 increased EcR-Usp interaction relative to the control samples as seen by FRET after acceptor photobleaching (**Fig. 4C**). Together, these data support a model wherein ATF4 and Usp compete against each other for binding to the limited pool of EcR available in the larval fat body.

### ATF4 and EcR coregulate *brummer*

Our experiments revealed that the ATF4-EcR complex can co-regulate transcriptional activation of the ATF4 target *4E-BP.* Both *ATF4* and *4E-BP* have been implicated in lipid metabolism in the *Drosophila* fat body^6,68^, which led us to consider if there were other lipid metabolism genes that were co-regulated by the ATF4-EcR complex. We recently reported that ATF4 can also act as a transcriptional repressor of the ATG lipase *bmm* in the adult fat body^29^. Based on this observation, we asked whether transcriptional repression by ATF4 in the larval fat body is also dependent on ATF4-EcR interaction. To test this, we employed a reporter wherein GFP expression is driven by a *bmm* intronic enhancer region containing ATF4-binding sites, *bmm1p^WT^-GFP*^8^.

We had previously validated the specificity of the *bmm1p^WT^-GFP* by mutating ATF4 binding sites in the enhancer region, which resulted in de-repression of *GFP* in S2 cells^8^. We generated transgenic flies containing the *bmm1p^WT^-GFP* reporter to examine the effects of EcR on ATF4-mediated *bmm* regulation. We were unable to detect GFP expression by microscopy in the fat body, indicative of very low levels of reporter expression. However, we were able to compare reporter expression by measuring *GFP* mRNA by RT-qPCR in isolated fat bodies. We found that depleting *EcR* resulted in increased *GFP* mRNA to a similar extent as that observed with *ATF4* knockdown (**Fig. 5A**). These data demonstrate that ATF4-mediated repression of *bmm* in the fat body is also mediated by the ATF4-EcR complex. Such a repressive role is also supported by our RNAseq analysis that revealed de-repression of another ATF4 repression target, *trbl*, with both *ATF4^RNAi^* and *EcR^RNAi^* (**Fig. 2E**). It is worth noting here that in adults, we previously reported that total *bmm* transcript levels were increased when *ATF4* was depleted in the fat body^8^. However, our RNAseq analysis of the larval fat body did not reveal a significant change in total *bmm* transcript levels with either *ATF4* or *EcR* knockdown (**Fig. S2**). This could be attributed to the presence of multiple *bmm* isoforms, only some which are regulated by ATF4-EcR, which our RNAseq analyses cannot parse. Alternatively, while the *bmm1p^WT^-GFP* is illustrative of ATF4-EcR-mediated transcriptional repression of a relatively small regulatory region of the *bmm* locus, ATF4 and EcR may not be the only transcription factors involved in *bmm* regulation in the third instar larval fat body. We tested this second possibility by examining changes in bmm protein levels.

**Fig 5.**
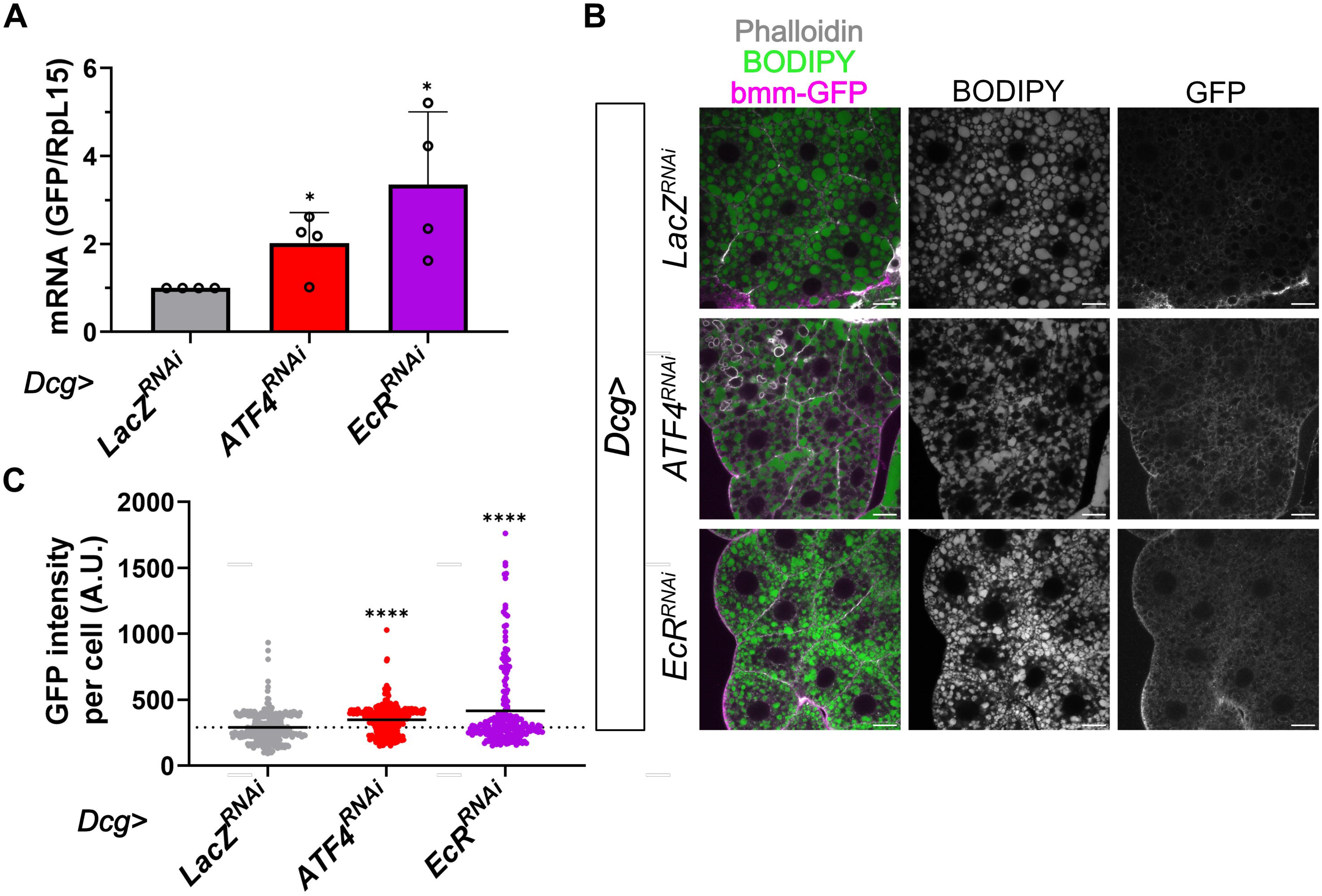
The ATF4-EcR complex represses *bmm* in the larval fat body. **A)** Analysis of bmm1p^WT^-GFP reporter expression as measured by RT-qPCR of GFP mRNA levels in isolated fat bodies also expressing the indicated RNAi lines driven by *Dcg-GAL4.* Data represent average of four biological replicates and error bars represent standard error of mean. **B)** Representative confocal images showing bmm-GFP expression (magenta) in fat bodies also expressing indicated RNAi lines driven by *Dcg-GAL4*. DAPI stains DNA (gray) and BODIPY stains lipid droplets (green). **C)** Quantification of bmm-GFP signal in individual adipocytes from **A**. Data represent mean from at least 18 animals collected from three independent crosses. The dotted line indicates the average of the control (*LacZ^RNAi^*) sample. Scale bar=25 μm. ****=p<0.00001; ***=p<0.0001; **=p<0.001; *=p<0.05. Only significant comparisons are shown. Statistical tests are described in the methods.

We examined expression of an endogenously tagged Bmm-GFP fusion protein^69^ in wandering third instar fat bodies from *Dcg>ATF4^RNAi^* animals and found that loss of *ATF4* resulted in a small but statistically significant increase in Bmm-GFP expression (**Fig. 5B-C**). Interestingly, *Dcg>ATF4^RNAi^* adipocytes also showed abnormal lipid droplet morphology as visualized with the neutral lipid dye BODIPY (**Fig. 5B**). Similarly, we found that *Dcg>EcR^RNAi^* fat bodies also showed a small increase in bmm-GFP expression and notably abnormal lipid droplet morphology (**Fig. 5B-C**). Together, these data indicate that ATF4-EcR mediated repression has small effects on Bmm protein levels, indicating other regulatory mechanisms may be in play at the *bmm* locus. However, the notable changes in lipid droplet morphology observed with both *Dcg>ATF4^RNAi^* and *Dcg>EcR^RNAi^* suggest a role for ATF4-EcR-mediated regulation of lipid metabolism possibly mediated by other transcriptional targets.

### EcR acts as a cofactor of ATF4 during homeostasis but not during methionine depletion

Our data thus far examined homeostatic ATF4-EcR interaction, prompting us to test whether EcR acts as a cofactor for ATF4 under conditions of exogenous stress. To do so, we employed a model of genetic nutrient deprivation wherein methionine is depleted in the fat via methioninase expression^70^. Methioninase is a bacterial enzyme that metabolizes methionine, resulting in amino acid depletion^70^. We previously showed that methioninase (abbreviated as met’ase in figures) expression in the larval fat activates ISR signaling^30^. We corroborated this by examining *4E-BP^intron^-DsRed* expression in the fat following methioninase expression (**Fig. 6A, B** column 1*, Dcg>LacZ^RNAi^*) and by quantifying *4E-BP* mRNA levels via qPCR (**Fig. S3**). We next asked whether such elevated ATF4 activity due to methionine depletion stress is EcR-dependent.

**Fig. 6.**
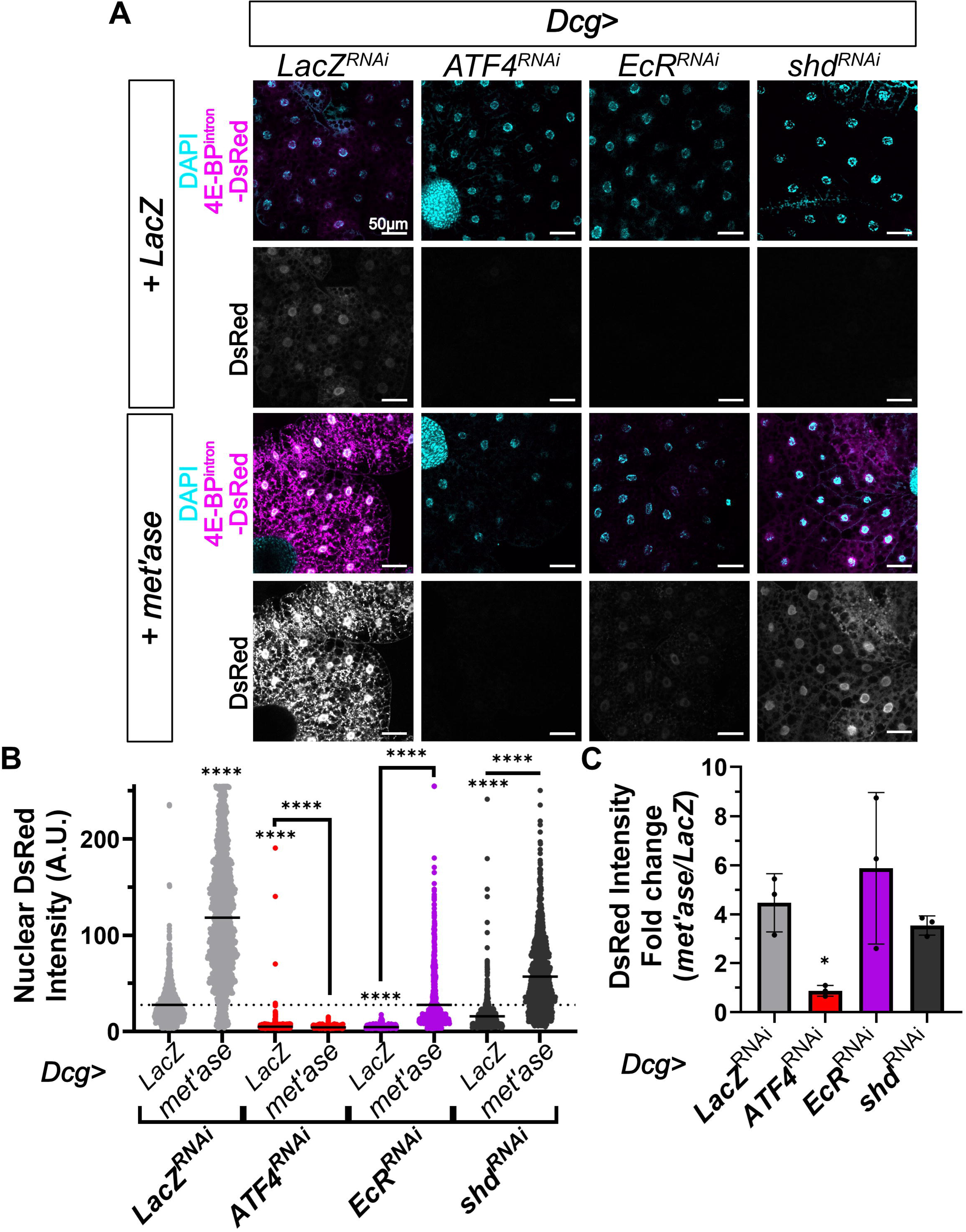
EcR is dispensable for increased ATF4 activity induced by exogenous stress. **A)** Representative confocal images showing 4E-BP^intron^-DsRed expression (magenta) in fat bodies where *Dcg-GAL4* drives either *UAS-methioninase (met’ase)* or a control *UAS-LacZ* expression. *Dcg-GAL4* simultaneously drives the expression of the indicated *UAS-RNAi* as well. **B)** Quantification of nuclear DsRed in individual adipocytes from **A**. Data represent mean from at least 18 animals collected from three independent crosses. The dotted line indicates the average of the control (*LacZ^RNAi^, LacZ*) sample. **C)** Fold change analysis of nuclear DsRed from **B.** Data represent the average of the mean induction across three independent experiments. Error bars represent standard error of mean. Scale bar=50 μm. ****=p<0.00001; ***=p<0.0001; **=p<0.001; *=p<0.05. Only significant comparisons are shown. Statistical tests are described in the methods.

As expected, depletion of *ATF4* resulted in diminished DsRed expression both in control *LacZ* and *methioninase* expressing fat bodies (**Fig. 6A-B**, *Dcg>LacZ, ATF4^RNAi^ vs Dcg>met’ase, ATF4^RNAi^*). In accordance with our initial observations, depleting *EcR* in the fat body resulted in lower *4E-BP^intron^-DsRed* expression in comparison to control animals (**Fig. 6A-B**, *Dcg>LacZ, LacZ^RNAi^ vs Dcg>LacZ, EcR^RNAi^,*). Strikingly though, we observed an increase in DsRed induction with methioninase in *EcR^RNAi^* animals (**Fig. 6A-B**, *Dcg>LacZ, EcR^RNAi^ vs Dcg>met’ase, EcR^RNAi^,*) which was further highlighted when the data were visualized as a fold induction (**Fig. 6C**). These results suggested that while EcR is required for ATF4 activity during homeostasis, it may be dispensable during stress. We further tested this indication by reducing intracellular 20E levels in the fat via *shd* depletion in animals expressing *LacZ* or *methioninase*. As with *Dcg>EcR^RNAi^*, while depletion of *shd* resulted in lower basal DsRed expression, it did not impact robust methioninase-induced increase in DsRed (**Fig. 6A-C**). Together, these results support a signaling paradigm where ATF4 and EcR interact in homeostatic adipocytes in a 20E-dependent manner and nutrient deprivation stress likely disrupts such interaction such that ATF4 activity is now independent of EcR.

## Discussion

Multiple studies have described ATF4 as a necessary factor in the development and homeostatic functioning of these same tissues ^1,7,9,28,29^, and yet other studies describe deleterious roles for aberrantly high ATF4 in metabolically active tissues in humans such as adipose and liver in disease states^10–12,15,71^. One prevailing hypothesis for how ATF4 executes these very different outcomes is due to its ability to bind to multiple bZIP partners^1^. Despite several such partners having been posited from biochemical and cell culture-based approaches, how ATF4 interaction with partners operates during homeostasis and changes during stress in metabolically active tissues has remained an open question. Our study tackles this problem by demonstrating stress-specific changes in the transcriptional activity of the ATF4-EcR complex using the *Drosophila* fat body. To our knowledge, this is the first *in vivo* demonstration of how ATF4 interaction with a partner is altered between homeostasis and stress conditions within the same tissue.

### Transcriptional regulation by the ATF4-EcR complex

Our transcriptomic and reporter-based analyses indicate that EcR can modulate a subset of ATF4-responsive genes, whereas reciprocal regulation appears far more limited, with ATF4 exerting little detectable influence on EcR transcriptional activity (**Figs. 1-2**). Interpretation of EcR-dependent effects on ATF4, however, is constrained by the relatively small number of well-defined ATF4 target genes in *Drosophila*, in contrast to the extensive catalog of EcR-regulated loci. As a result, the conclusion that ATF4 does not broadly regulate EcR targets is better supported by our transcriptomics data (**Fig. 2C**), although we did observe notable exceptions to this trend (**Fig. 2D**). To investigate these exceptions, we used publicly available modENCODE data^72,73^ to examine ATF4 occupancy on Hr4, Eip74EF, and Sox14. While we found a few potential binding sites in the Hr4 intronic region for ATF4 (**Fig. S4A**), none were recorded in Eip74EF and Sox14. These analyses leave open the possibility that there may be select EcR targets such as Hr4 that are coregulated by the ATF4-EcR complex.

Clustering of transcription factor binding sites within a regulatory region has been shown to increase regulatory capacity by enabling cooperative or additive transcriptional control^74^. Occupancy analysis based on modENCODE data show that EcR binds both the *bona-fide* ATF4 targets, *4E-BP* and *bmm* (**Fig. S4A**). Further analysis using the known JASPAR position weight matrix for EcR for the *4E-BP* intronic enhancer sequence, where there are several validated ATF4 binding sites^48^, showed the presence of multiple EcR consensus regions proximal to ATF4 binding sites (**Fig. S4B, Table S3**). This was also true for the *bmm* enhancer region where we had previously identified and validated ATF4 binding sites^8^ (**Table S3**). These clustered binding sites for ATF4 and EcR are likely to strengthen ATF4-EcR interaction, which we were able to detect even whilst using ATF4-GFP (**Fig. 1D**) which is encoded by a hypomorphic allele^32^. It is notable here that Alphafold3 analysis^75^ also supports direct interaction between the N-terminal domain of EcR-B2 and the ATF4 bZIP domain (**Fig. S4C**), similar to what was demonstrated by *in vitro* binding experiments^34^.

The increased ATF4-EcR signaling observed upon *usp* depletion (**Fig. 4**) suggests that EcR availability may be a limiting factor for engaging regulatory elements in the fat body. At loci with clusters of ATF4 and EcR binding sites such as the *4E-BP* intronic enhancer, the regulatory potential may be shaped not only by motif density and arrangement but also by the local concentration and availability of transcriptional partners. Indeed, clusters of low-affinity sites have been shown to reach high occupancies and support gene activation *in vivo* at physiologically relevant transcription factor concentrations^76^, indicating that factor availability interacts with cis-regulatory architecture to determine regulatory output. This model also aligns with the requirement for 20E in ATF4-EcR-mediated transcriptional activation of the *4E-BP* intronic enhancer (**Fig. 3**), as 20E is likewise necessary for EcR-Usp-mediated transcriptional activation.

### Regulation of lipid metabolism by the ATF4-EcR complex

Both ATF4 and EcR have independently been implicated in lipid metabolic regulation in the *Drosophila* fat body^77–79^, suggesting that these factors participate in overlapping metabolic control pathways. Consistent with this, the two loci identified here as ATF4-EcR targets, *4E-BP* and *bmm*, have each been previously linked to lipid metabolism, albeit primarily in the context of the adult fat body. 4E-BP has been characterized as a “metabolic brake” that limits lipid breakdown^68^, likely through translational control of metabolic effectors, whereas bmm encodes a triglyceride lipase that directly promotes lipid mobilization^80^. While both 4E-BP and bmm contribute to lipid homeostasis, their functions have been most extensively examined under nutrient deprivation or catabolic conditions^48,68,80,81^, leaving their regulation and roles in anabolic contexts less well understood. Given this, it is important to acknowledge here that transcriptional regulation is inherently context dependent, and our findings may therefore reflect features specific to the third instar larval fat body, a tissue that exists in a largely anabolic state. However, it is worth noting that the metabolic profile of this tissue has been likened to that of proliferating cancer cells^82–84^, making our findings here relevant in other tissues in anabolic states. This notion is further supported by a demonstrated role for ATF4 in driving overproliferation in other *Drosophila* tissues modeling cancer-like metabolic states^85^.

Both cancer cells and the developing third instar larval fat body prioritize the conversion of glucose into lactate even when oxygen is plentiful^83^. This shift is not merely for ATP production but primarily to generate the carbon-based precursors (amino acids, lipids, and nucleotides) required for rapid biomass accumulation^82^. In this context, repression of *bmm* by ATF4-EcR can be readily interpreted as a mechanism to minimize lipid breakdown and favor lipid accumulation. Consistent with this model, loss of *ATF4* results in reduction in overall lipid content accompanied by abnormal lipid droplet morphology^6,8^, which is phenocopied with loss of EcR (**Fig. 5**). Notably, we had previously reported anomalous lipid droplet morphology with loss of *ATF4* in the adult fat body^67^, and vertebrate ATF4 is similarly thought to repress *ATGL*^86^, the vertebrate ortholog of bmm, highlighting the conserved nature of ATF4-mediated regulation of lipolytic pathways. This conservation raises the question of the corresponding nuclear receptor partner for ATF4 in vertebrates. The closest vertebrate ortholog of EcR is the liver X receptor (LXR)^87–89^, a type II nuclear receptor with well-established roles in lipid metabolism^90^. Our work lays a conceptual foundation for exploring whether ATF4-LXR interactions contribute to lipid metabolic regulation in vertebrate systems, and more broadly suggests that ATF4 may engage distinct nuclear receptor partners in a context-dependent manner to coordinate metabolic gene expression.

### EcR-independent ATF4 signaling during nutrient deprivation stress

Under conditions of exogenous stress, such as our genetic nutrient deprivation model, we find that loss of EcR does not impair further induction of ATF4-mediated transcriptional activation (**Fig. 6**). This observation suggests that EcR is not required for amplification of canonical ATF4 transcriptional responses during stress, although it does not exclude additional ATF4-independent roles for EcR in this context. Indeed, both increased 20E levels and enhanced EcR activity in response to nutrient deprivation and other stressors have been robustly documented^91–94^, indicating that EcR participates broadly in the organismal stress response. An important caveat of our analysis is that it focuses specifically on ATF4-EcR mediated transcriptional activation during stress using the *4E-BP^intron^-DsRed* reporter, rather than repression. Thus, our study does not address potential transcriptional repression regulatory modes that may be differentially engaged during stress. In this regard, EcR-Usp-mediated transcriptional repression has been proposed to occur in a ligand-independent manner^95–98^, raising the possibility that EcR contributes to stress-responsive gene regulation through mechanisms not captured by our reporter. One intriguing model is that ATF4 may switch from acting as a repressor of *bmm* under homeostatic conditions to an activator during nutrient deprivation, with this functional transition being mediated by 20E-dependent changes in EcR activity or complex composition. Such a model could reconcile the apparent dispensability of EcR for stress-induced ATF4 activation with the well-established role of ecdysone signaling in starvation and developmental stress responses.

It is also important to consider that ATF4 may pair with another interaction partner(s) both during homeostasis and stress. *Drosophila* ATF4 is capable of partnering with multiple bZIP transcription factors^35^, and it may be that distinct interaction partners predominate under different physiological conditions. Considering the clustered binding site architecture observed at ATF4-EcR-regulated loci, it is plausible that ATF4 transcriptional output during stress is shaped by cooperative interactions among several transcription factors, with EcR representing only one of multiple potential contributors. Defining how these factors dynamically assemble at shared regulatory elements, and how their interactions are modulated by stress cues, will be an important area for future investigation.

## Materials and Methods

### Drosophila fly lines used

Animals were raised on ‘R’ food formulation obtained from LabExpress, Inc (by weight: 80% distilled water, 1% agar, 8% molasses, 7% corn meal, 4% yeast, 0.5% propionic acid, 0.25% methyl p-hydroxybenzoate). Fly stocks were maintained at room temperature and experimental crosses were reared at 25°C under light/dark conditions. All stocks used in this study are listed in **Table 1** and genotypes for each experiment are listed in **Table 2**. All experiments were performed on fat bodies dissected from female wandering third instar larvae collected from timed 4-hour egg lays in uncrowded vials, except for **Fig. S1A-B** which shows data from male larvae.

**Table 1.** List of fly stocks.

| **Fly strain** | **Stock number/reference** |
| --- | --- |
| Dcg-GAL4 | BDSC_7011 |
| UAS-LacZ<sup>RNAi</sup> | Kang et al. 2016<sup>48</sup> |
| UAS-ATF4<sup>RNAi</sup> | VDRC109014 |
| UAS-EcR<sup>RNAi</sup> | BDSC_58286 |
| UAS-EcR<sup>RNAi</sup> (#2) | BDSC_29374 |
| 4E-BP<sup>intron</sup>-DsRed | Kang et al. 2016<sup>48</sup> |
| UAS-shd<sup>RNAi</sup> | BDSC_67356 |
| UAS-EcI<sup>RNAi</sup> | Okamoto et al, 2018<sup>105</sup> |
| UAS-usp<sup>RNAi</sup> | BDSC_27258 |
| 4E-BP<sup>intron</sup>-GFP | Grmai et al. 2024<sup>30</sup>. |
| UAS-EcR B2<sup>F645A</sup> | BDSC_9450 |
| UAS-EcR-ΔN | BDSC_6868 |
| w<sup>1118</sup> | BDSC_3605 |
| crc<sup>GFSTF</sup> (a.k.a. ATF4 -GFP) | BDSC_59608 |
| usp-GFP | BDSC_38672 |
| bmm-GFP | BDSC_94600 |
| bmm1p<sup>WT</sup>-GFP | Generated in this study |
| UAS-LacZ | BDSC_3956 |
| UAS-methioninase | Parkhitko et al, 2021<sup>106</sup> |

**Table 2.**
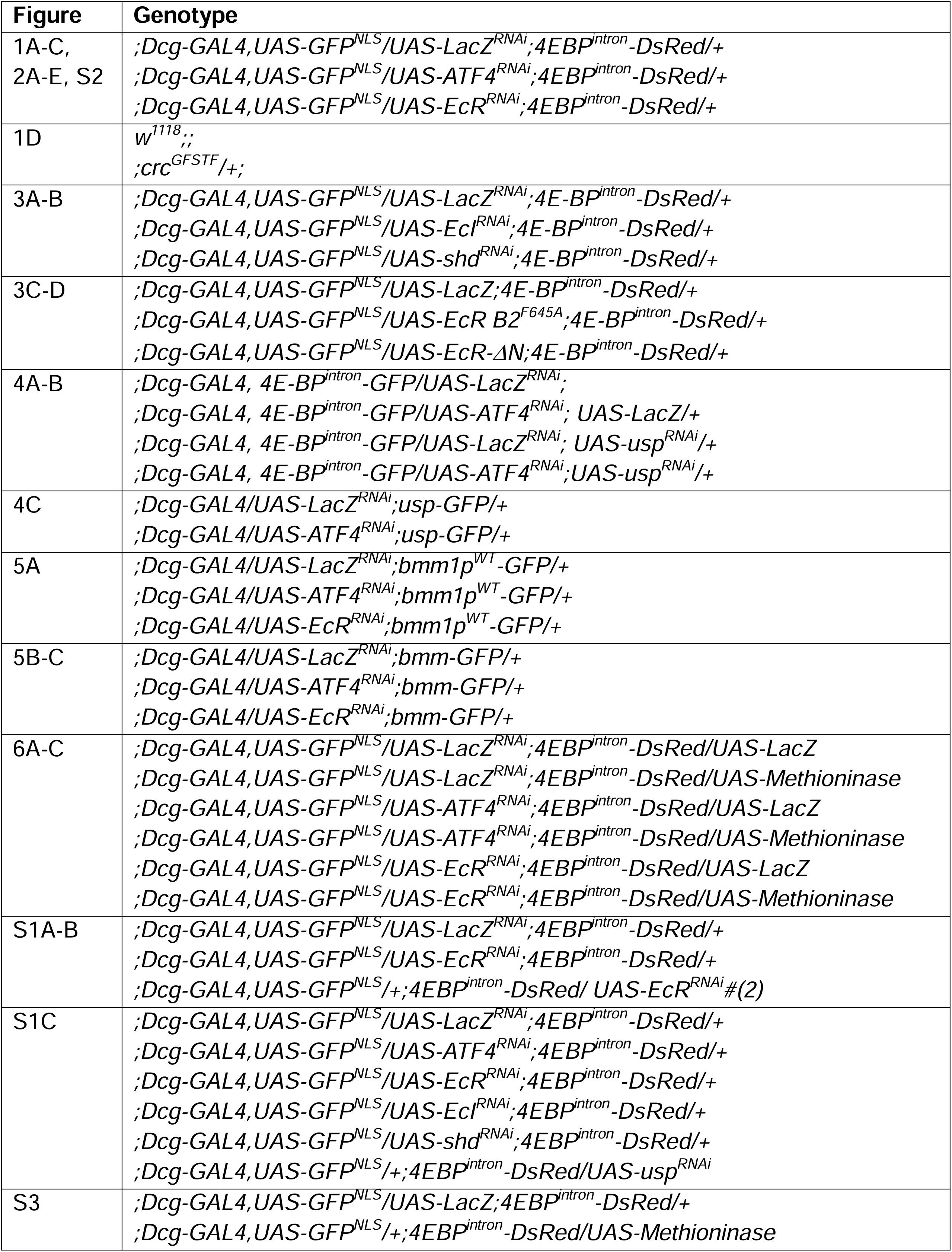
List of genotypes by figure.

### Immunofluorescence and quantification

Fat bodies were dissected from wandering third instar larvae and fixed in phosphate buffered saline (PBS), 4% PFA, 0.1% Tween-20 for 20 minutes with rocking at room temperature. Samples were washed twice in PBS for 5 minutes. Samples were incubated overnight at 4°C in chicken α-GFP antibody (1:250; Thermo Fisher Scientific #501962090) diluted in PBS. Samples were incubated for 1 hour in the dark in donkey α-chicken Alexa Fluor 647 (1:500; Thermo Fisher Scientific #GFP-101AP) diluted in PBS. Samples were kept in the dark from here on. Samples were incubated for 30 minutes in Rhodamine phallodin (1:500, Thermo Fisher Scientific #50646256) and boron-dipyrromethane 493/503 (BODIPY) (1:250, Thermo Fisher Scientific #D3922). Samples were incubated for 5 minutes in PBS containing 4′,6-diamidino-2-phenylindole (DAPI) (1nM, Thermo Fisher Scientific #574810) which was followed by two washes in PBS. Samples were mounted and imaged using a Nikon A1 confocal microscope using a 20x air lens for *4E-BP^intron^* reporters and 60x oil immersion lens for bmm-GFP experiments.

All image analyses were performed using FIJI ver. 2.3 on a single slice of a confocal image. Nuclear fluorescent protein intensity for individual adipocytes expressing 4E-BP^intron^-DsRed or - GFP were measured by generating a DAPI-based mask to delineate nuclear area. The same mask was used to calculate nuclear area of individual adipocytes. For bmm-GFP expression, boundaries for individual cells were hand-marked and GFP intensity was calculated.

### Quantitative RT-qPCR

Fat bodies were dissected from four wandering third instar larvae from 4-hour egg lays. RNA was extracted from these fat bodies using TRIzol (Thermo Fisher Scientific #15596026) and cDNA was made using Maxima H minus RT (Thermo Fisher Scientific #EP0751). Quantitative RT-qPCR was performed using qMAX^TM^ Green qPCR mix (MIDSCI SKU PR2001-H-1000) and measured on a C1000 Touch^TM^ Thermal Cycler. Primers used in this study are listed in **Table 3**.

**Table 3.**
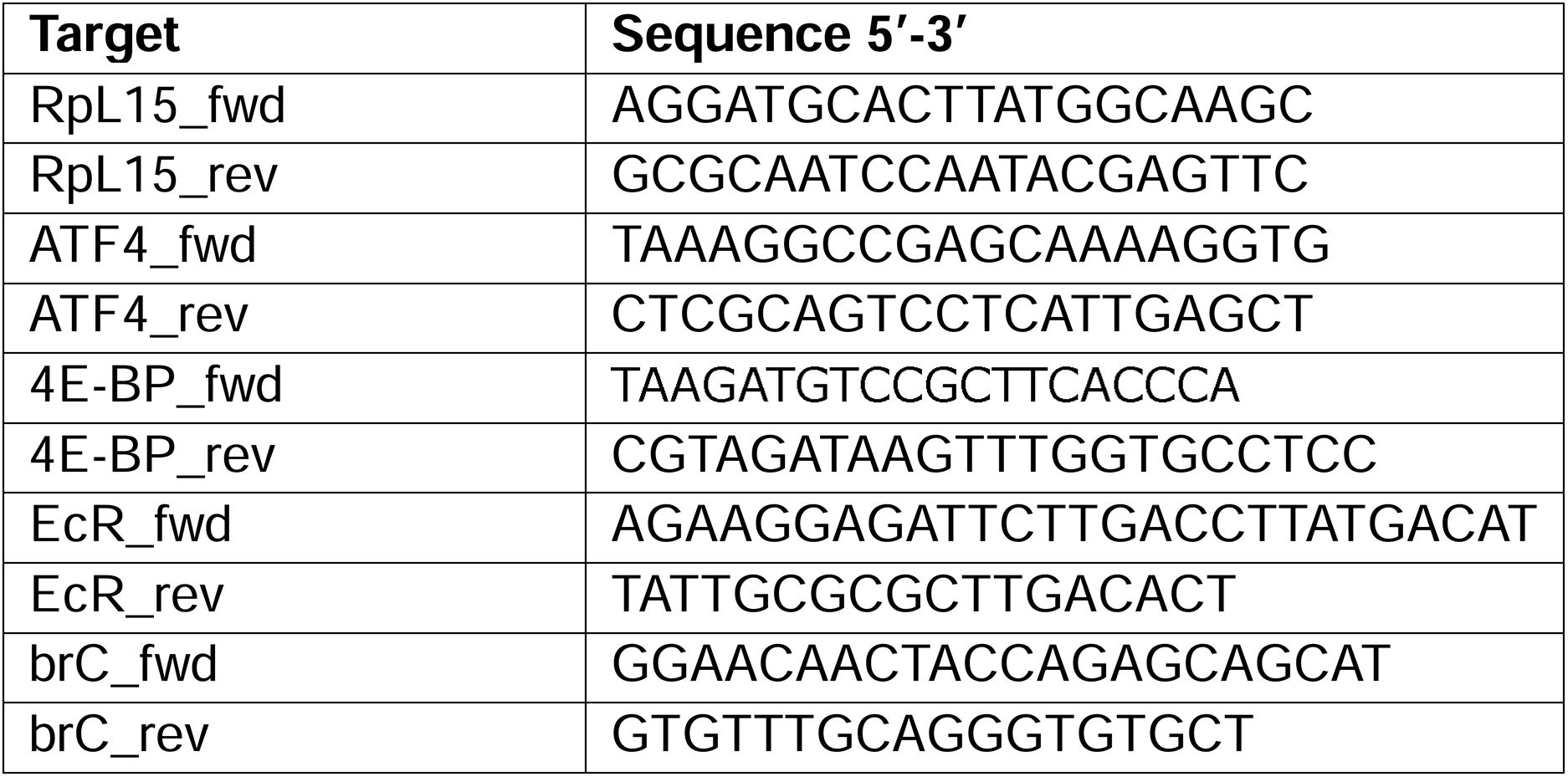
List of RT-qPCR primers.

### RNAseq

Fat bodies were collected from four wandering third instar larva and RNA was extracted as described above for RT-qPCR. Duplicate RNA samples from independent crosses were submitted for each genotype to the Plasmidsaurus RNAseq analysis pipeline. In brief, the pipeline utilizes a poly-T bead-based extraction of total RNA, followed by cDNA preparation, second-strand synthesis, tagmentation, library indexing and amplification. Samples are then Illumina sequenced using a 3’ end counting approach, with an average read length of 90 bps.

FastQ files are generated with BCL Convert v4.3.6 and fqtk v0.3.1. Reads are filtered using FastP v0.24.0 to ensure poly-X tail trimming, 3’ quality-based tail trimming, a minimum Phred quality score of 15, and a minimum length requirement of 50 bp. The sequences were aligned to the BDGP6.54_62 (dm6) reference genome using STAR aligner v2.7.11 to generate BAM files, which were coordinate sorted using samtools v1.22.1. PCR and optical duplicates were removed using UMICollapse v1.1.0 and gene expression was quantified using featureCounts (subread package v2.1.1) with strand-specific counting, multi-mapping read fractional assignment, exons and 3’ UTR as the feature identifiers. Finally, differential expression was analyzed using edgeR v4.0.16 with filtering for low-expressed genes with edgeR::filterByExpr with default values. These analyzed data are available in **Table S1, S2** and presented as differential expression analyses in **Fig. 2, S2**.

### FRET analysis and quantification

Dissected fat bodies were fixed in PBS+4% PFA+0.1% Tween-20 for twenty minutes with rocking at room temperature. Samples were washed twice in PBS for 5min. Samples were incubated overnight at 4°C in mouse α-EcR common antibody (1:50, DSHB Ag10.2) and chicken α-GFP antibody (1:250, Thermo Fisher Scientific #501962090) diluted in PBS. Samples were incubated for 1 hour in the dark in donkey α-mouse Cy3 antibody (1:1000, Jackson ImmunoResearch #715-165-150) and donkey α-chicken Cy5 antibody (1:1000, Jackson ImmunoResearch #703-175-155) diluted in PBS. Samples were kept in the dark from here on. Samples were incubated in DAPI diluted in PBS as described above for 5 min followed by two washes in PBS and subsequently mounted in Gelvatol mounting media. Prebleach and postbleach images were captured after photobleaching Cy5 antibody with 638nm laser for 45 seconds. Cy3 channel was collected in 30 channel-bins from 569nm to 748nm. The area under the curve for nuclear Cy3 was calculated from spectral channels 569.64-634.50nm to measure the relative change in FRET. Confocal images were captured using a Nikon A1 Spectral confocal microscope with 60x oil immersion lens.

### Generation of *bmm1p^WT^-GFP* transgenic flies

The plasmid encoding the bmm1p^WT^-GFP reporter was cloned previously^99^. The plasmid was injected into strain 8622 by BestGene Inc, targeting it to the attP2 landing site. Successful transformants were selected using the w+ marker.

### modENCODE and binding site analyses

ATF4- and EcR-occupancy analyses at various loci were examined using the modENCODE data accession ENCFF252CAI and ENCFF162EYM respectively. Occupancy was visualized and graphed using IGV^100^.

For binding site analyses, the position weight matrix for *Drosophila* ATF4 (crc) and EcR were obtained from the JASPAR database^101^ (motif IDs MA2208.1 and MA2223.1 respectively). The motifs were analyzed using the FIMO tool in MEME suite^102^ against enhancer element sequences. The sequence for the enhancer elements analyzed from *4E-BP* and *bmm* were previously described^48,99^ and are provided in **Table S3**.

### Statistical analyses

The figure legends describe the sample sizes for the corresponding data. For all fluorescence image quantifications with multiple comparisons (**Figs. 1C, 3B, 3D, 4D, 5C, 6A, 6B, S1B, S1C**), significance was tested using a Kruskal-Wallis test with Dunn’s correction for multiple comparisons. Image quantifications with paired comparisons (**Figs. 1D, 4C**) were made using the Mann-Whitney test for nonparametric data in all instances of FRET data when we could not assume Gaussian distribution. For RT-qPCR data where control samples were normalized to 1 (**Figs.1A, 5A, S3**), where each data point has a corresponding measurement across all conditions, when making multiple comparisons we utilized a Friedman test for nonparametric data with Dunn’s correction for multiple comparisons. RNA-seq data report p-values using the Benjamini-Hochberg method which controls for false discovery rate to determine significant gene expression.

## Contributions and funding

D.V. conceived the project. M.M., L.G., and D.V. designed the experiments. M.M. and D.V. performed the experiments, analyzed the data, and drafted the initial manuscript. M.M., L.G., and D.V. revised the manuscript. M.M. and D.V. are supported by NIH R35GM150516 (to D.V.) and a teaching fellowship from the University of Pittsburgh School of Medicine (to M.M.). L.G. is funded by NIH R00GM149982 (to L.G).

## Acknowledgements

We would like to thank publicly available model organism resources that fueled our research: FlyBase, Bloomington Drosophila stock center, and Vienna Drosophila stock center. We thank Dr. Claudette St. Croix and the Center for Biologic Imaging at the University of Pittsburgh for extensive support with FRET imaging data acquisition and analysis. We would like to thank Dr. Naoki Yamanaka for sharing transgenic fly lines to deplete EcI. We would like to thank all members of our lab for discussion and feedback on the project, and Dr. Lesley Weaver for critical reading of the manuscript.

Stocks obtained from the Bloomington Drosophila Stock Center (NIH P40OD018537) and Vienna Drosophila stock center^103^ were used in this study. We used FlyBase^104^ (release FB2024_01) to identify phenotypes and stocks in this study.

---

**Table S1 and S2.**

Differential gene expression analysis and counts per million from RNAseq analysis of *LacZ^RNAi^* vs. *ATF4^RNAi^* (S1) or vs. *EcR^RNAi^* (S2).

**Table S3.**

List of binding sites predicted by FIMO analysis (see Methods) of the intronic enhancers from the *4E-BP* and *bmm* locus using the known position matrix of ATF4 and EcR from JASPAR. The value of a motif occurrence is defined as the probability of a random sequence of the same length as the motif matching that position of the sequence with as good or better a score. The score for the match of a position in a sequence to a motif is computed by summing the appropriate entries from each column of the position-dependent scoring matrix that represents the motif. The q-value of a motif occurrence is defined as the false discovery rate if the occurrence is accepted as significant.

**Fig. S1.**
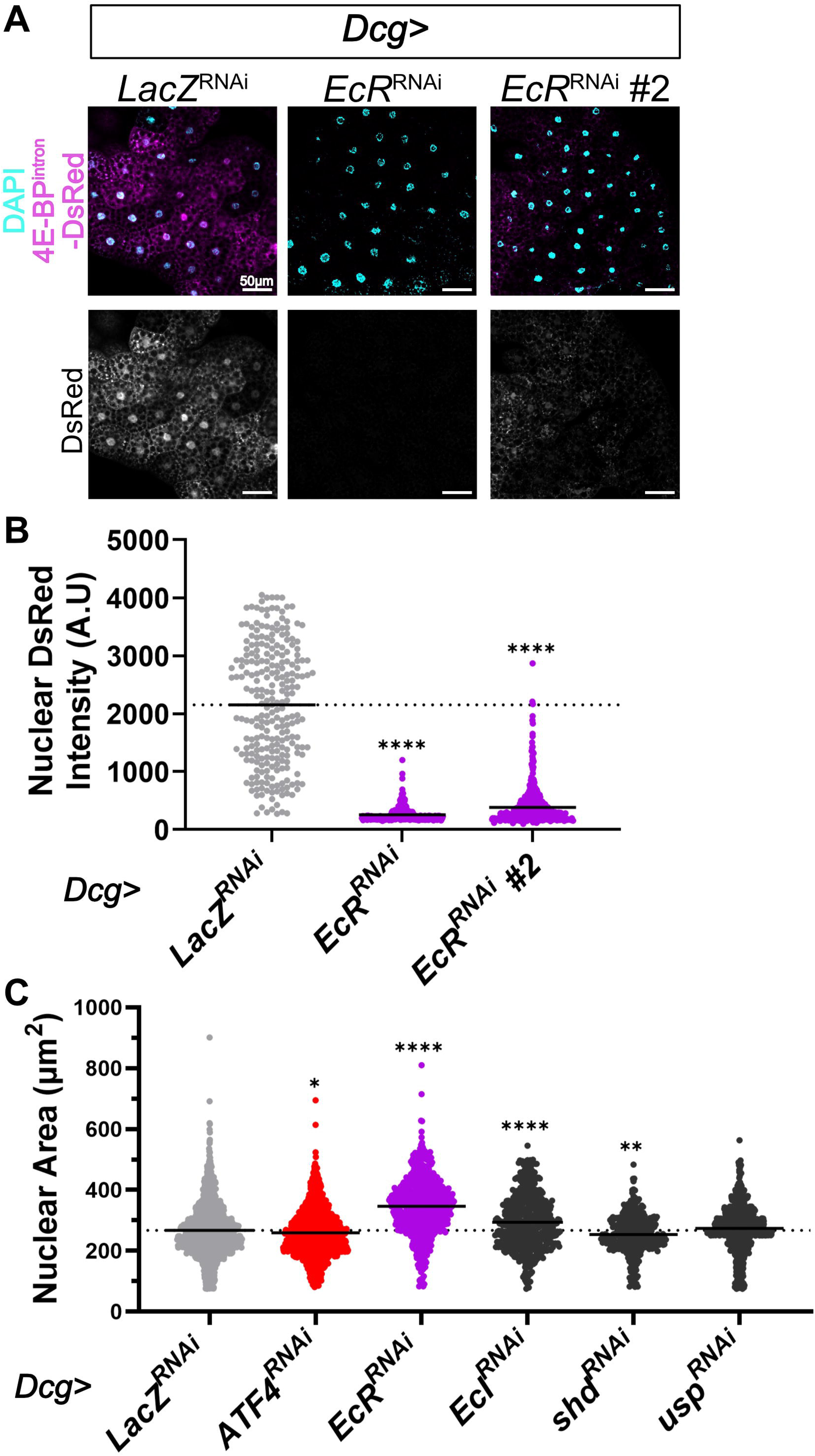
EcR-mediated decrease in ATF4 signaling is not sexually dimorphic or due to developmental timing. **A)** Representative confocal images showing 4E-BP^intron^-DsRed reporter expression (magenta) in male fat bodies also expressing indicated RNAi lines driven by *Dcg-GAL4*. Nuclei are counterstained with DAPI (cyan). **B)** Quantification of nuclear DsRed in individual adipocytes from **A**. Data represent mean from at least 10 animals collected from two independent crosses. The dotted line indicates the average of the control (*LacZ^RNAi^*) sample. **C)** Quantification of nuclear area in individual adipocytes from Fig. 1B, 3A, and 4A. The dotted line indicates the average of the control (*LacZ^RNAi^*) sample. Scale bar=50 μm. ****=p<0.00001; ***=p<0.0001; **=p<0.001; *=p<0.05. Only significant comparisons are shown. Statistical tests are described in the methods.

**Fig. S2.**
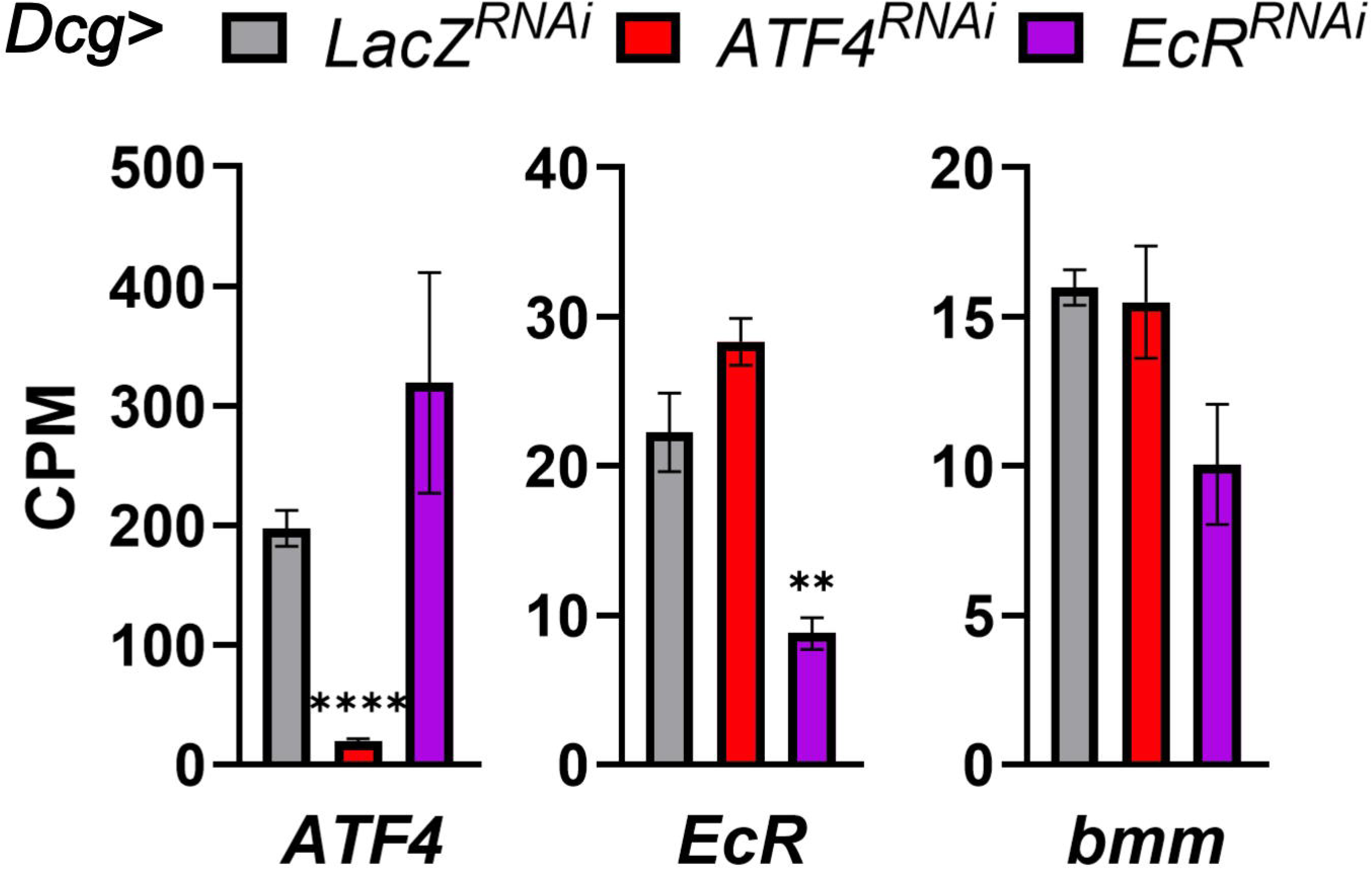
Total bmm transcript levels are unchanged with *ATF4* or *EcR* depletion. Counts per million (CPM) for the indicated transcripts from the RNAseq analysis in Fig. 2. ****=p<0.00001; ***=p<0.0001; **=p<0.001; *=p<0.05. Only significant comparisons are shown. Statistical tests are described in the methods.

**Fig. S3.**
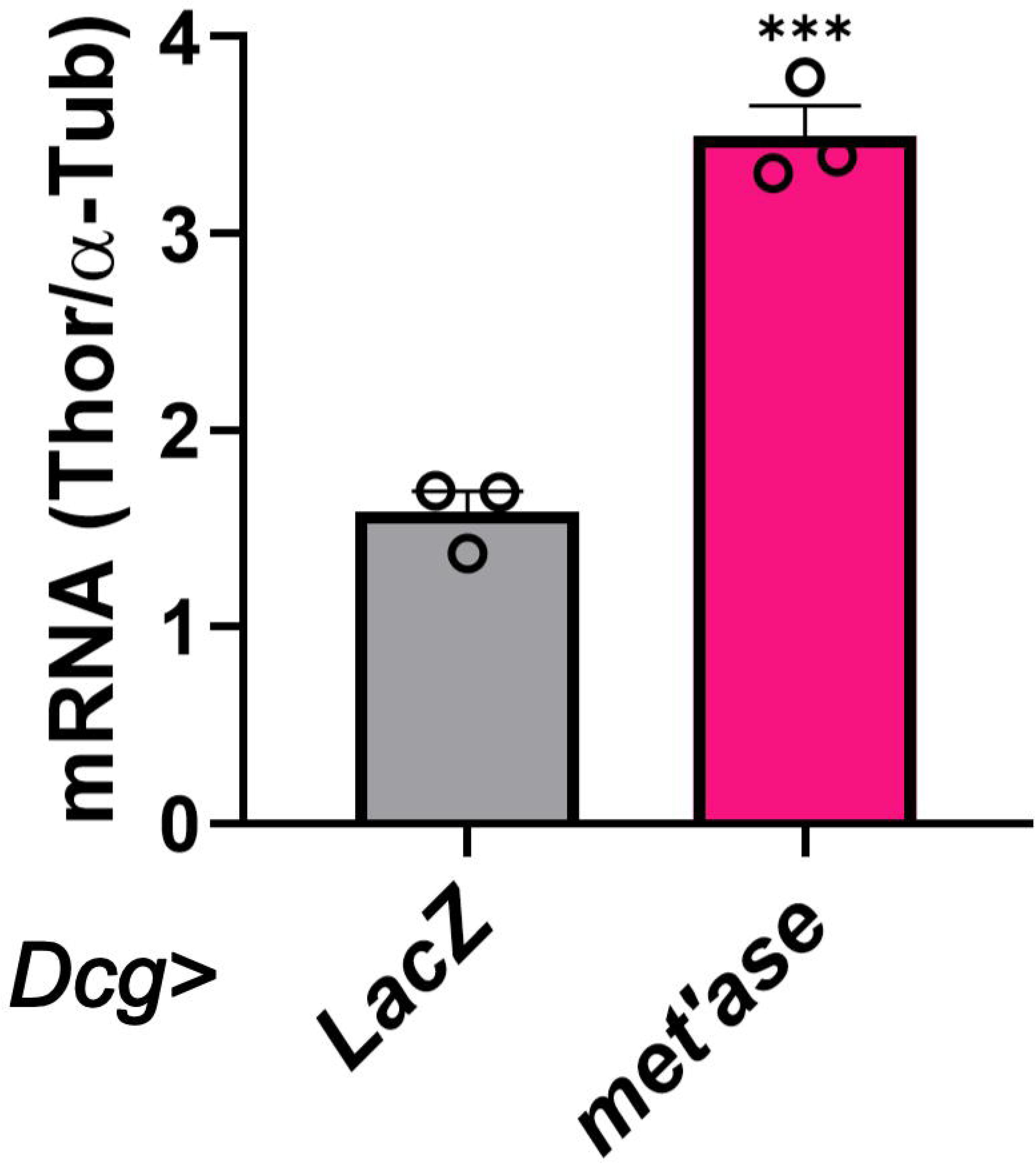
Methioninase expression further induces ATF4 activity in the fat body. RT-qPCR analysis of *4E-BP* mRNA expression in fat bodies from **A.** ****=p<0.00001; ***=p<0.0001; **=p<0.001; *=p<0.05. Only significant comparisons are shown. Statistical tests are described in the methods.

**Fig. S4.**
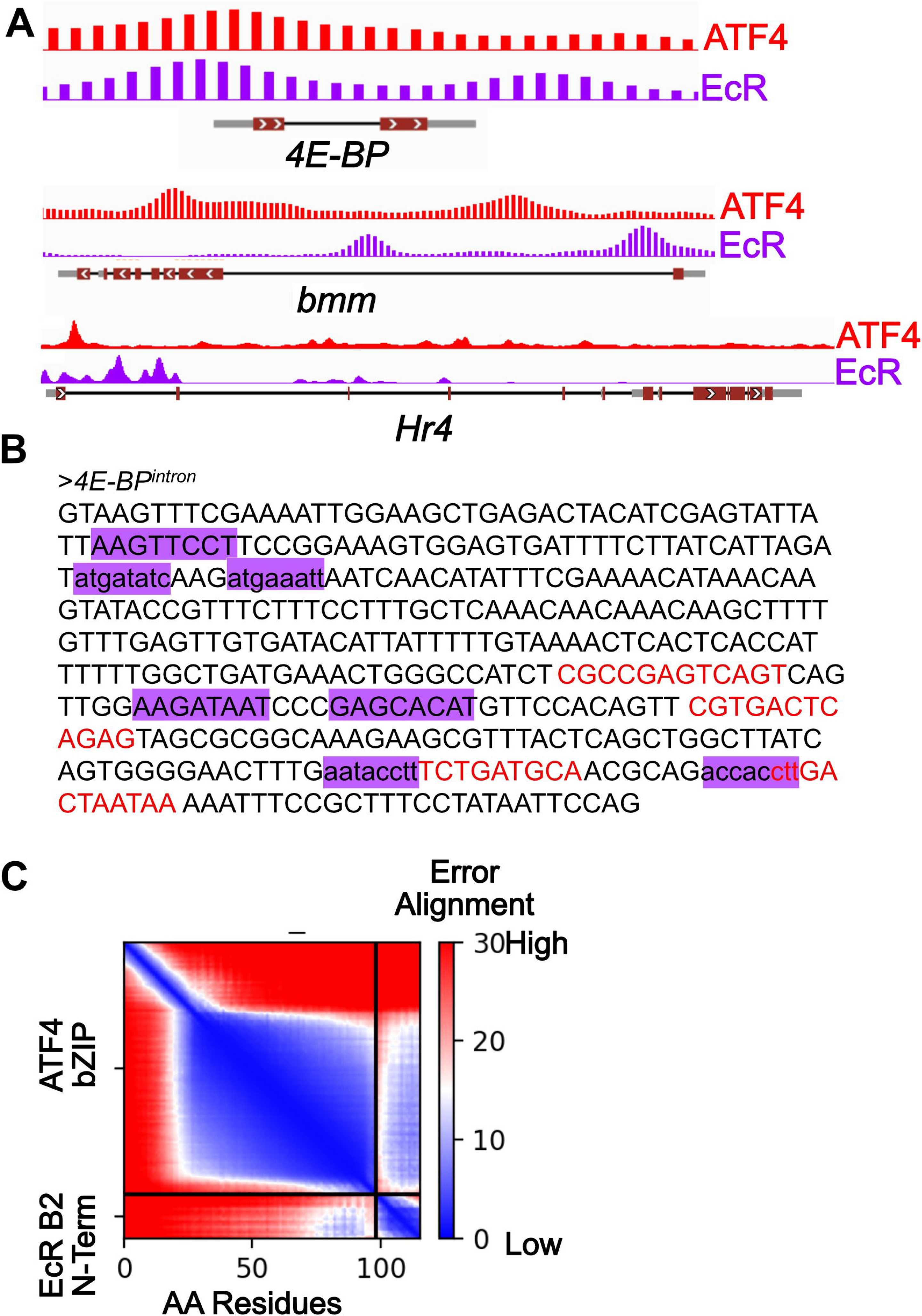
ATF4 transcriptional targets contain EcR-binding sites. **B)** Analysis of modENCODE data showing occupancy of ATF4 and EcR on the indicated genetic loci. **C)** Decorated sequence of the intronic enhancer of the *4E-BP* locus showing ATF4 (red) and EcR (orange) binding sites as predicted by the known position weight matrices for these factors (see Methods). **D)** The AlphaFold Predicted Aligned Error (PAE) between ATF4 basic leucine zipper and EcR-B2 N-terminal region. PAE measures confidence in the relative distance and orientation between residues in the indicated proteins, low PAE (blues) indicates high confidence in the special relationship and conversely high PAE (reds) indicate low confidence.

